# What’s the matter with MICs: The contribution of nutrients and limiting resources to the pharmacodynamics of antibiotics and bacteria

**DOI:** 10.1101/2022.09.30.510422

**Authors:** Brandon A. Berryhill, Teresa Gil-Gil, Joshua A. Manuel, Andrew P. Smith, Ellie Margollis, Fernando Baquero, Bruce R. Levin

## Abstract

The minimum inhibitory concentration (MIC) of an antibiotic required to prevent replication is used both as a measure of the susceptibility/resistance of bacteria to that drug and as the single pharmacodynamic parameter for the rational design of antibiotic treatment regimes. MICs are estimated *in vitro* under conditions optimal for the action of the antibiotic. However, bacteria rarely grow in these optimal conditions. Using a mathematical model of the pharmacodynamics of antibiotics, we make predictions about the nutrient dependency of bacterial growth in the presence of antibiotics. We test these predictions with experiments in a rich media and a glucose-limited minimal media with *Escherichia coli* and eight different antibiotics. Our experiments uncover properties that question the sufficiency of using MICs and simple pharmacodynamic functions as measures of the pharmacodynamics of antibiotics under the nutritional conditions of infected tissues. To an extent that varies among drugs: (i) The estimated MICs obtained in rich media are greater than those estimated in minimal media. (ii) Exposure to these drugs increases the time before logarithmic growth starts, their lag. (iii) The stationary phase density of *E. coli* populations declines with greater sub-MIC antibiotic concentrations. We postulate a mechanism to account for the relationship between the sub-MIC concentration of antibiotics and the stationary phase density of bacteria and provide evidence in support of this hypothesis. We discuss the implications of these results to our understanding of the MIC as the unique pharmacodynamic parameter used to design protocols for antibiotic treatment.

## Introduction

Fundamental to the rational, as opposed to purely empirical, design of antibiotic treatment regimens are the bacterial-dependent pharmacodynamics, PD [1–3], and host-dependent pharmacokinetics, PK [1, 2, 4], and their relationship known as PK/PD indices. The PKs of antibiotics are measured from the changes in concentrations of the drug in the serum of treated hosts, often utilizing uninfected volunteers [5, 6]. Although different measures of PK of the antibiotic are employed for PK/PD indices [7], most commonly a single measure of the PD of the antibiotic and its target bacteria are employed. This measure is the minimum concentration of the antibiotic necessary to prevent the replication of the bacteria, the <u>M</u>inimum <u>I</u>nhibitory <u>C</u>oncentration [8].

MICs are measured in vitro, most commonly by two-times serial dilution [9, 10]. In this method, bacteria are inoculated into a rich media at a density of 5×10^5^ cells per mL and the antibiotic is added at a high concentration to a single culture. The cultures are serially diluted by a factor of two and incubated for a defined period of time. At this time, the optical densities of the antibiotic-exposed cultures are estimated by eye, looking for the well where turbidity drastically decreases [11]. The antibiotic concentration at which this sharp decrease in turbidity is seen is called the MIC. This method tends to have a lower throughput than what is needed for clinical laboratories, thus other standardized, machine-based protocols are used for estimating MICs. Though, these methods such as the VITEK [12, 13] are also based on optical density, and work by comparing the growth of a known isolate to an unknown isolate. Non-optical density-based methods do exist, such as E-Tests [14, 15].

It has been shown that the value of the MIC estimated for antibiotic and bacteria pairs are critically dependent on the density of bacteria exposed to the antibiotics when starting the microtiter broth dilution [16]. In this investigation we use mathematical models to explore the relationship between the richness of the media and concentration of a limiting resource, on the pharmacodynamics of antibiotics and bacteria. Using *Escherichia coli* growing in glucose-limited minimal medium and Lysogeny Broth (LB) we explore the fit of these models to the pharmacodynamics of antibiotics of eight different classes.

The results of our study question the sufficiency and validity of existing procedures to estimate MICs and using this parameter to provide a quantitatively accurate picture of the pharmacodynamics of antibiotics and bacteria [17]. We demonstrate that: i) The PD of antibiotics are critically dependent on the media used, (ii) exposure to sub-MIC concentrations of antibiotics increases the time before bacteria start to grow, (iii) exposure to sub-MIC concentrations of antibiotics decreases the growth rate of these bacteria in proportion to the concentration of the drug, (iv) the stationary phase density to which the bacteria grow is inversely related to the concentration of antibiotics, (v) sub-MIC concentrations of antibiotics increase the nutrient requirements of the treated bacteria. We discuss the implications of these PD results to the design and evaluation of antibiotic treatment protocols.

## Results

### A generalized model of the pharmacodynamics of antibiotics and bacteria and its predictions

For our model of the pharmacodynamics of antibiotics and bacteria we combine Hill functions [16] with a resource-limited growth model [18]. In accord with this extended model, the hourly rate of growth of the bacteria is defined with respect to the concentration of the antibiotic (A μg/mL) and the limiting resource (r μg/mL) as shown in Equation 1.

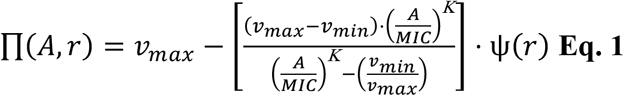

The parameter *v_MAX_* is the maximum rate of growth per cell per hour of the bacteria in the absence of the antibiotic, where *v_MAX_*>0. *v_MIN_* is the minimum growth rate per cell per hour of the bacteria in the presence of the antibiotic, *v_MIN_*<0. *MIC* is the minimum inhibitory concentration in μg/mL of the antibiotic. *K* is a shape parameter that determines the acuteness of the π(A,r) function, the greater the value of *K* the more acute. Ψ(*r*), defined in Equation 2, expresses the relationship between the rate of growth of the bacteria and the concentration of the limiting resource, r. The parameter k in μg/mL is the Monod constant which is the concentration of the resource when the growth rate in the absence of antibiotics is half of its maximum value, *v_MAX_*.

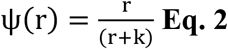

To explore the relationship between antibiotic concentration, limiting resource concentration, and bacterial density, we present three coupled differential equations based on the Hill functions.

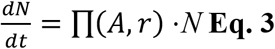

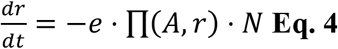

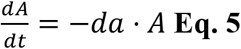

∏*(A,r)* is the Hill function designated rate of growth or death of the bacteria when the concentration of the antibiotic is A and the resource is r (Eq. 3). The conversion efficiency, e μg, is the amount of the limiting resource needed to produce a new cell (Eq. 4). The parameter da is the hourly rate of decline in the concentration of the antibiotic (Eq. 5).

In Figure 1A and 1B we illustrate the predicted relationship between the concentration of the antibiotic and the rates of growth and death of bacteria in an environment with an indefinite amount of resource, Ψ(*r*)=1.0, for a bacteriostatic (Conditon I), weakly bactericidal (Condition II), and strongly bactericidal (Conditon III) antibiotic based on different Hill function parameters. Other than indicating the minimum concentration of the antibiotic that prevents growth, the MIC of an antibiotic provides no information about the dynamics, rate of growth, or rate of death of the bacteria. Antibiotics with the same MIC can be bacteriostatic (Figure 1A Condition I) or they can be bactericidal (Figure 1A Condition II and Condition III). Moreover, the rate of growth of bacteria exposed to sub-MIC concentrations of bacteriostatic antibiotics (Figure 1B condition I) would be greater than bacteria exposed to highly bactericidal antibiotics (Figure 1B condition III).

**Figure 1.**
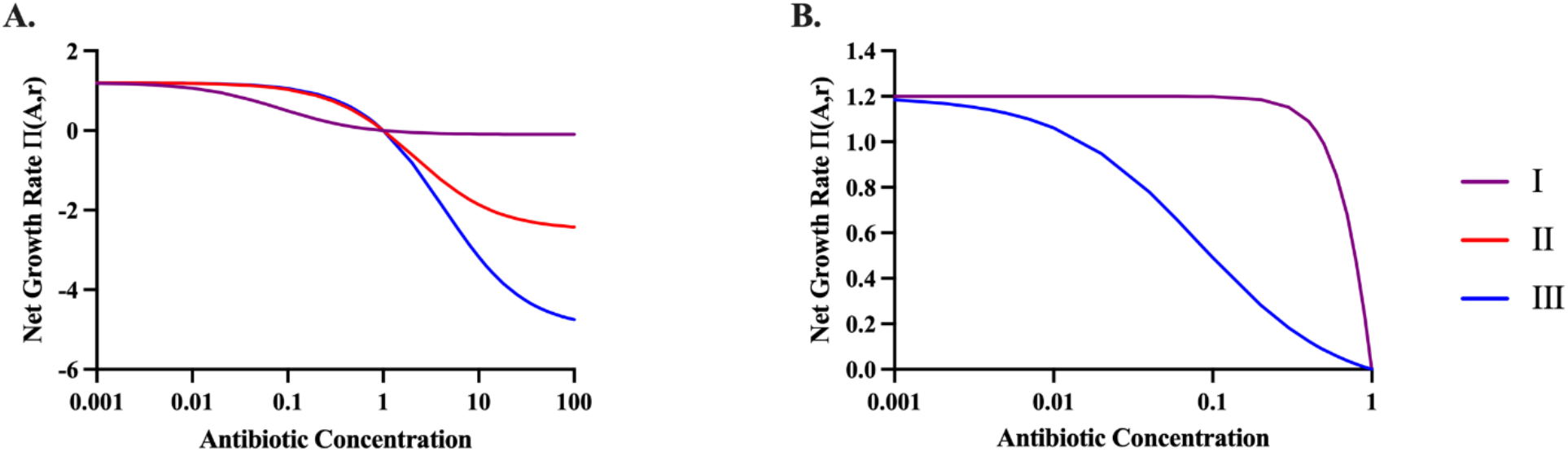
Predicted growth rates of antibiotic-exposed bacteria. To illustrate the properties of this model, we explore three different parameters sets I, II, and III. Condition I represents a bacteriostatic antibiotic (*v_MIN_* = −0.0001). Condition II represents a weakly bactericidal antibiotic (*v_MIN_* = −2.5). Condition III represents a strong bactericidal antibiotic (*v_MIN_* = −5.0). Parameters used for these simulations are e= 5×10^−7^ μg/per cell, *v_MAX_* = 1.2 per cell per hour, K=1, k=1, MIC=1.0. **(A)** and **(B)** The relationship between the concentration of an antibiotic and the rates of growth and death of bacteria *π*(*A*, *r*) anticipated by the pharmacodynamic function for the described values of the Hill function parameters (Equation 1). In these figures the population is not limited by the resource Ψ(*r*)=1.0. **(A)** All concentrations, **(B)** Sub-MIC concentrations of the antibiotic.

In Figure 2 we follow theoretical changes in the density of bacteria exposed to different concentrations of a strongly bactericidal antibiotic as predicted by our model (Equations 1–5). Under these conditions, the model predicts that the density achieved by the bacterial population is the same at all concentrations of the antibiotic below MIC. Stated another way, the theory predicts that the reduction in growth rate due to the presence of the antibiotic only increases the time before the population reaches stationary phase.

**Figure 2.**
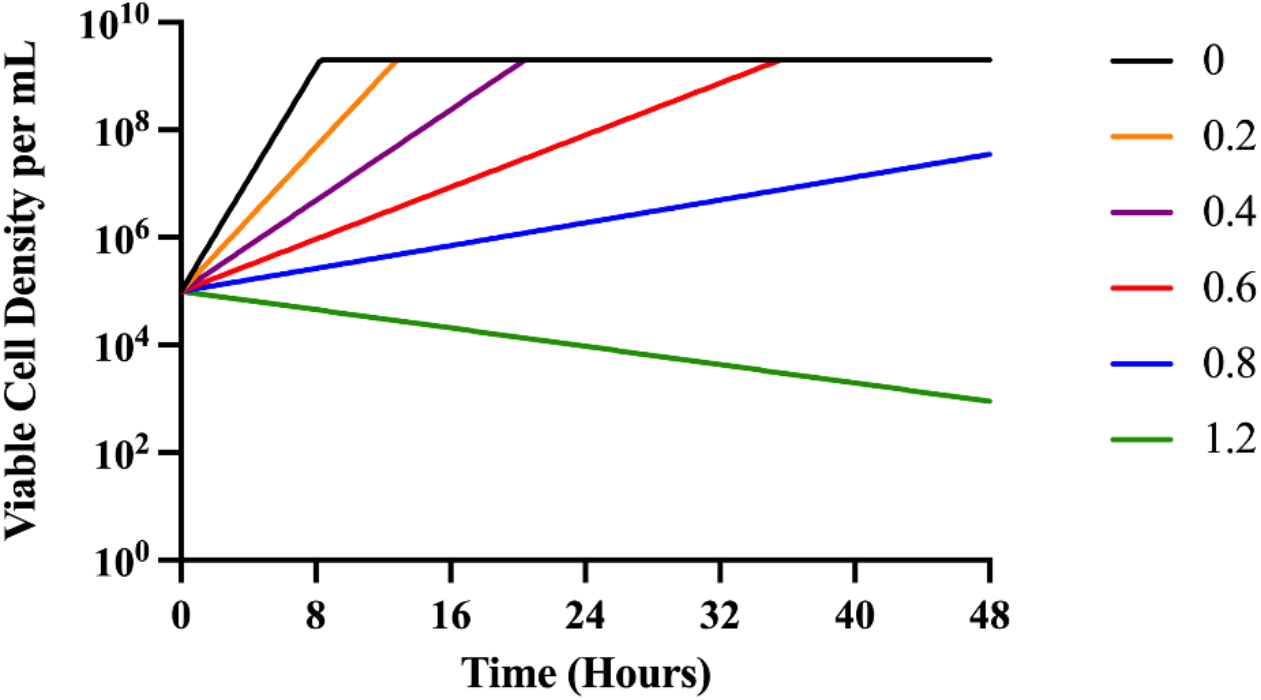
Predicted changes in density of antibiotic-exposed bacteria. To illustrate the properties of this model, we explore only the strong bactericidal antibiotic condition (Condition III). Condition III represents a strong bactericidal antibiotic with *v_MIN_* = −5.0. Parameters used for these simulations are e=5×10^−7^ μg/per cell, *v_MAX_* =1.2 per cell per hour, K=1, k=1, MIC=1.0. Changes in the viable density of bacteria anticipated from the Hill function model for different concentrations of the antibiotic, respectively 0 to 1.2 times MIC. In these numerical solutions to equations 2–4, we assume the concentration of the limiting resource (r) is 1000 when the bacteria are first introduced to the antibiotic and the concentration of the antibiotic does not decay, da=0.

### Experimental exploration of MIC as the primary PD parameter

#### Minimum Inhibitory Concentrations

We open our experimental exploration of the pharmacodynamics of antibiotics and bacteria with a consideration of the relationship between the estimated MICs of antibiotics with *E. coil* MG1655 growing in LB and Davis Minimal (minimal) medium with 1000μg/mL glucose (Table 1).

**Table 1.** Estimated MICs of *E. coli* MG1655 for eight antibiotics in LB and glucose-limited minimal medium.

| Antibiotic | LB MIC (μg/mL) | Minimal MIC (μg/mL) |
| --- | --- | --- |
| AZM | 25 | 6.25 |
| CIP | 0.03 | 0.03 |
| TET | 2.5 | 0.8 |
| FOF | 37.5 | 25 |
| GEN | 12 | 0.75 |
| RIF | 25 | 12.5 |
| CHL | 6.25 | 6.25 |
| CRO | 0.05 | 0.03 |

For ciprofloxacin, fosfomycin, chloramphenicol, rifampin, and ceftriaxone the MICs estimated in broth and minimal medium are equal or nearly so. For azithromycin, gentamicin, and tetracycline the MICs estimated in broth are substantially greater than that in glucose-limited minimal medium. In supplemental Table 1, we show the MICs estimated with minimal medium with different carbon sources. Notably, the MICs of certain drugs such as azithromycin, fosfomycin, and ceftriaxone are lower in alternative carbon sources than they are in glucose. The values presented for the MICs of these drugs were estimated as the first concentration at which the optical density declined substantially and approached that of the media. As shown in supplemental figure S1, sub-MIC concentrations in most antibiotics lead to a decrease in the optical density in proportion to the drug concentration, an effect we call MIC ooze.

#### Growth dynamics at sub-MIC concentrations of eight antibiotics in LB and minimal media

In Figure 3 and Figure 4 we follow the changes in optical density of *E. coli* MG1655 exposed to different antibiotics in LB and minimal media, respectively. A detailed comparison of the parameters estimated from this analysis (maximum growth rate, length of the lag phase, and maximum optical density) are available in Supplemental figures Fig S2 and Fig S5, Fig S3 and Fig S6, and Fig S4 and Fig S7, respectively.

**Figure 3.**
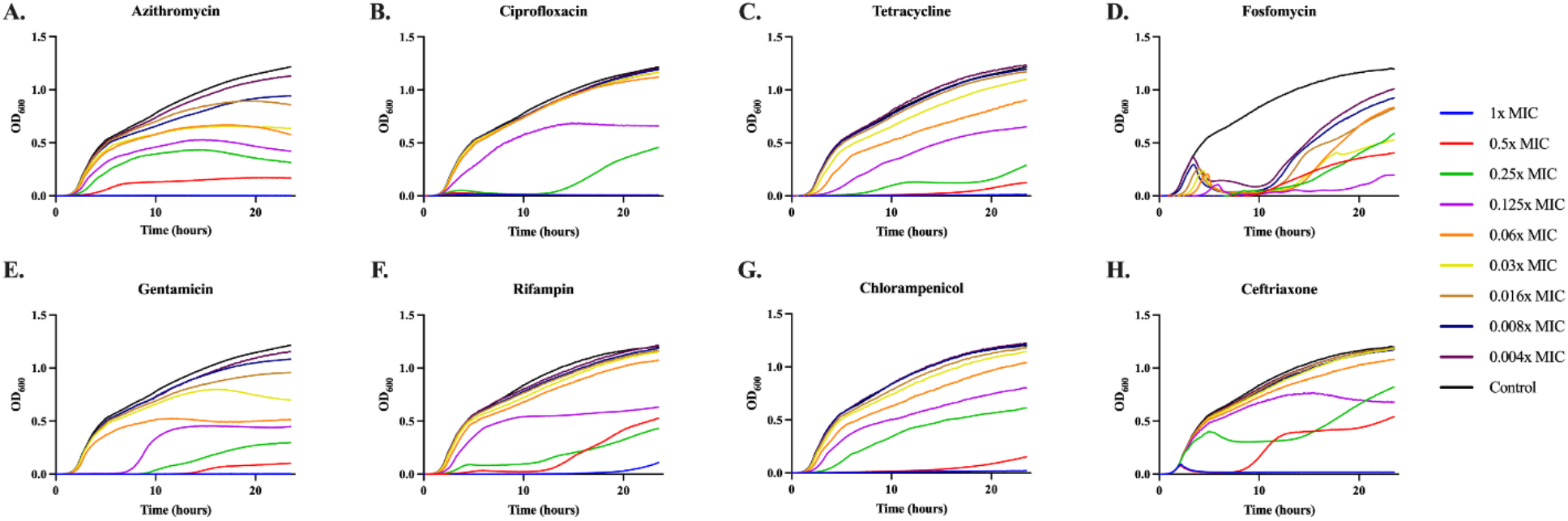
Growth dynamics in LB. Changes in optical density (600nm) of *E. coli* MG1655 exposed to different sub-MIC concentrations of eight antibiotics for 24 hours in LB. Lines are representative of the average of five technical replicas and normalized to the time zero optical density. Each concentration is shown as a fraction of the MIC for the noted drug (Table 1): 1x (blue), 0.5x (red), 0.25x (green), 0.125x (purple), 0.06x (orange), 0.03x (yellow), 0.016x (brown), 0.008x (dark blue), 0.004x (dark purple), and a no drug control (black).

**Figure 4.**
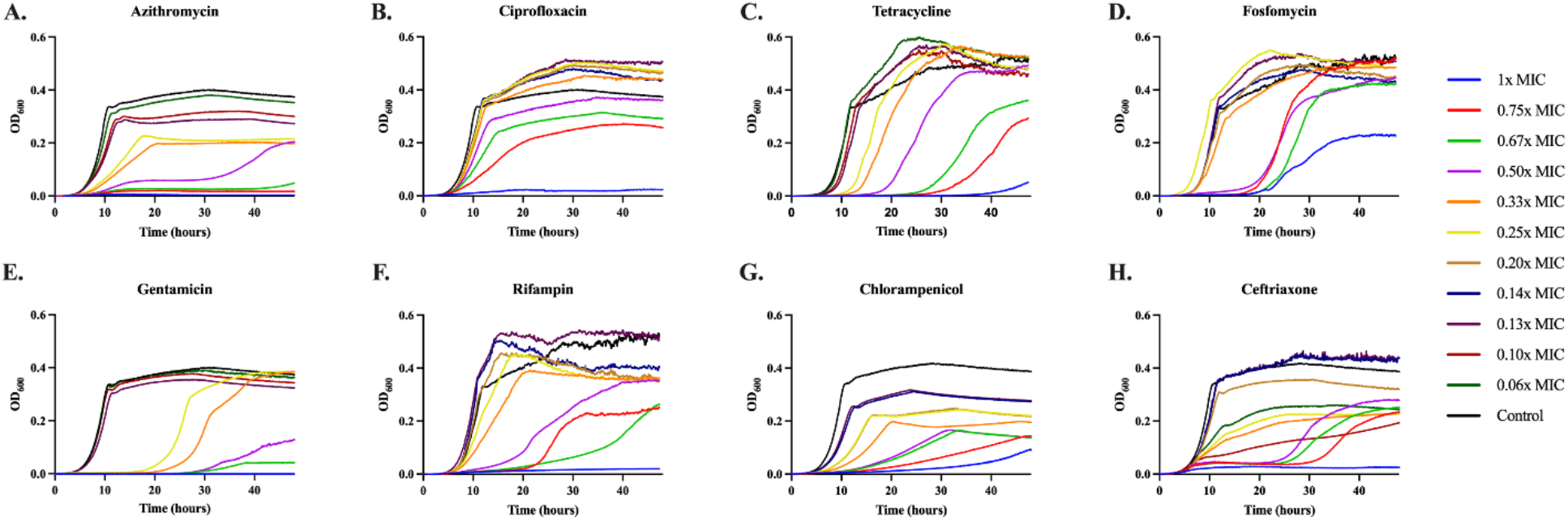
Growth dynamics in minimal medium. Changes in optical density (600nm) of *E. coli* MG1655 exposed to different sub-MIC concentrations of eight antibiotics for 48 hours in glucose-limited minimal medium. Lines are representative of the average of five technical replicas and normalized to the time zero optical density. Each concentration is shown as a fraction of the MIC for the noted drug (Table 1): 1x (blue), 0.75x (red), 0.67x (green), 0.50x (purple), 0.33x (orange), 0.25x (yellow), 0.20x (brown), 0.14x (dark blue), 0.13x (dark purple), 0.10x (dark red), 0.06x (dark green), and a no drug control (black).

The dynamics of change in optical densities of MG1655 vary among the drugs in both broth and minimal medium. For all eight antibiotics, these dynamic data indicate concentration-dependent variation in the time before the populations start to grow, the rate of growth, and the density achieved at 24 hours for LB and 48 hours for the minimal medium.

In Figure S2 and S5 we plot the relationship between the estimated maximum growth rates as a function of sub-MIC concentration of the antibiotics in LB (Fig S2) and minimal (Fig S5). With the exception of fosfomycin in both medias, the max rate of growth increases with the declining concentration of antibiotic. The observation that at higher concentrations the antibiotics would be reducing the growth rate of the bacteria, to an extent greater than at lower concentrations, is anticipated and central to the use of the pharmacodynamic Hill functions.

In Figure S3 and S6, we plot the amount of time required before the optical density exceeds 0.020, the relative lag for different concentrations of the antibiotics in LB and minimal medium, respectively. In both medias, there is a concentration dependent increase in the lag time, with higher concentrations of the drug leading to longer lag times.

The optical density at 24 hours in LB and 48 hours in minimal are presented in Figure S4 and S7, respectively. In both media, for all of the antibiotics the maximum optical density increases with the decline in the concentration of the antibiotic. This decrease in the maximum optical density is associated with a decrease in the viable colony forming units (CFU) of these cultures (Figure S8).

Consistently across all three parameters, fosfomycin, gentamicin, and rifampin have high variability between replicates. This high variability can be explained by the emergence of both genotypic (fosfomycin Figure S9 and rifampin Figure S10) and seemingly phenotypic (gentamicin) resistance. For fosfomycin and rifampin resistances emerge independently across replicas as shown in Figure S9 and S10. For fosfomycin an increase in MIC of over 11-fold was seen between pre- and post- antibiotic exposed cultures, while for rifampin an increase of 35 times was seen. This rapid selection for fosfomycin resistance is well known as it has been shown that there is a large inoculum density effect when performing MICs with this drug [19]. When MG1655 was exposed to gentamicin, a large number of small colony variants were recovered. One possibility is that the exposure to the drug during LB double dilution rapidly selects for a resistant minority population. As a test of this, we performed an E-Test on MG1655 with gentamicin and found the MIC to be 24 times lower by E-Test as compared to double dilution, a result we believe to be consistent with heteroresistance.

#### Consumption of a limited resource in sub-MIC treated cultures

The association between the concentration of the drug and the final optical density of each culture is striking and inconsistent with the predictions of Figure 2. In accord with the model, we would expect drug concentration to simply increase the amount of time needed to reach stationary phase, but not to impact the level of stationary phase. This is clearly not the case. One possibility is that when exposed to the drugs, the bacteria become less efficient in the consumption of resource. The utilization of glucose limited minimal media allows for direct measurement of the glucose level (Figure 5) and in accord with the above hypothesis, we would expect there to be no excess glucose after 48 hours of bacterial growth. Our results are consistent with this hypothesis. At super-MIC concentrations of the drugs, we see low to no consumption of the glucose, while at sub-MIC concentrations we see consumption of almost all the limiting resource. With the exception of ceftriaxone, which at super-MIC concentrations consumes a large amount of glucose.

**Figure 5.**
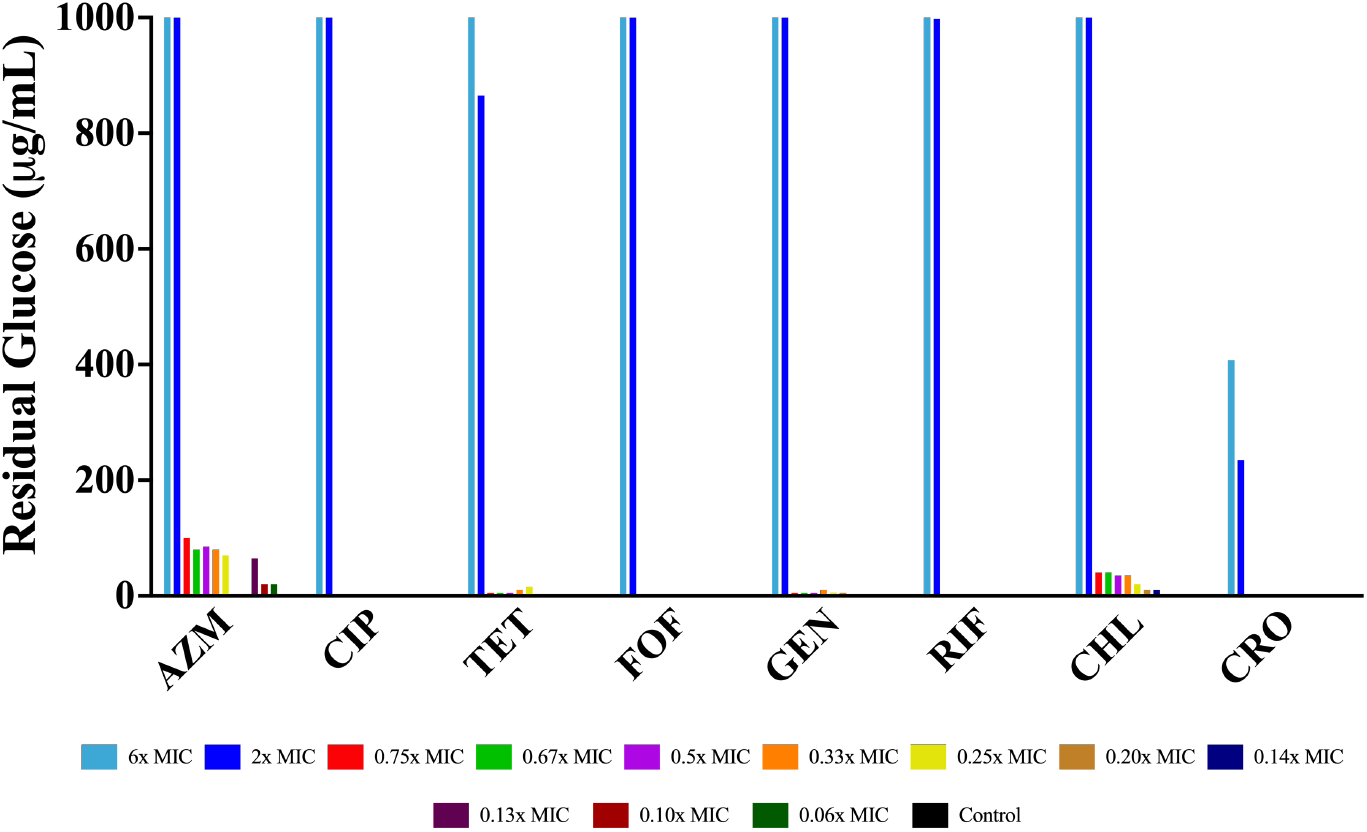
Final Glucose Concentrations. Amount of glucose (μg/mL) in each culture of *E. coli* MG1655 exposed to different super- and sub-MIC concentrations of eight antibiotics after 48 hours of incubation in glucose-limited minimal medium.

#### Antibiotic dependent resource consumption

In accord with Figure 2, the stationary phase densities of bacteria exposed to sub-MIC concentrations of antibiotics should be independent of the concentration of the drug. That is clearly not the case as the stationary phase density declines with the concentration of the antibiotic (Figures S4 and S7). One possible explanation is that in addition to reducing the rate of growth of the *E. coli*, the antibiotic increases the amount of resource necessary to produce a new cell [17]. As such, we have updated Equation 4 to make resource consumption dependent on the concentration of the antibiotic (Eq. 6). By modifying the resource uptake equation, we get:

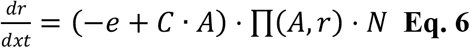

where C is a constant that determines the extent to which e is increased. In Figure 6 we illustrate the changes in density predicted by this updated model. As shown in Figure 3 and 4, sub-MIC concentrations of antibiotics are able to decrease the stationary phase density achieved.

**Figure 6.**
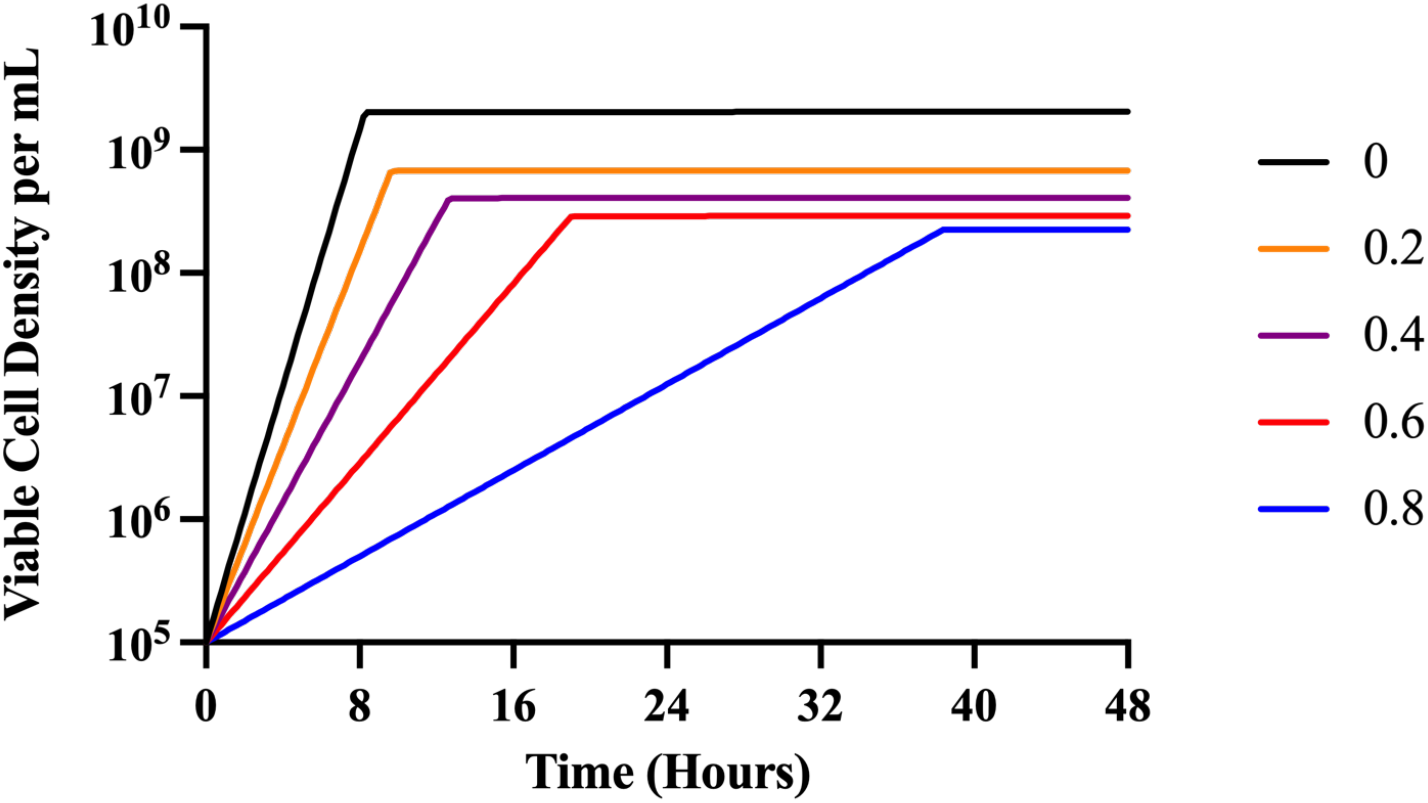
Predicted changes in density of antibiotic-exposed bacteria with antibiotic-dependent resource consumption. To illustrate the properties of this model, we explore only the strong bactericidal antibiotic condition (Condition III). Condition III represents a strong bactericidal antibiotic with *v_MIN_* = −5.0. Parameters used for these simulations are e=5×10^−7^ μg/per cell, *v_MAX_* =1.2 per cell per hour, K=1, k=1, MIC=1.0. Changes in the viable density of bacteria anticipated from the Hill function model for different concentrations of the antibiotic, respectively 0 to 0.8 times MIC. In these numerical solutions, we assume the concentration of the limiting resource (r) is 1000 when the bacteria are first introduced to the antibiotic and the concentration of the antibiotic does not decay, da=0, and that C= 5×10^−6^.

#### Rates of antibiotic-dependent growth and mortality

To explore the relationship between the concentration of the antibiotic and the rates of growth and death of *E. coli* MG1655, we performed time-kill experiments. Overnight cultures of the bacteria in LB (Figure 7) or minimal medium (Figure 8) were mixed with different concentrations of the antibiotics above and below the MIC estimated for the drugs in the noted media. Viable cell densities (CFUs) of the cultures were estimated at different times.

**Figure 7.**
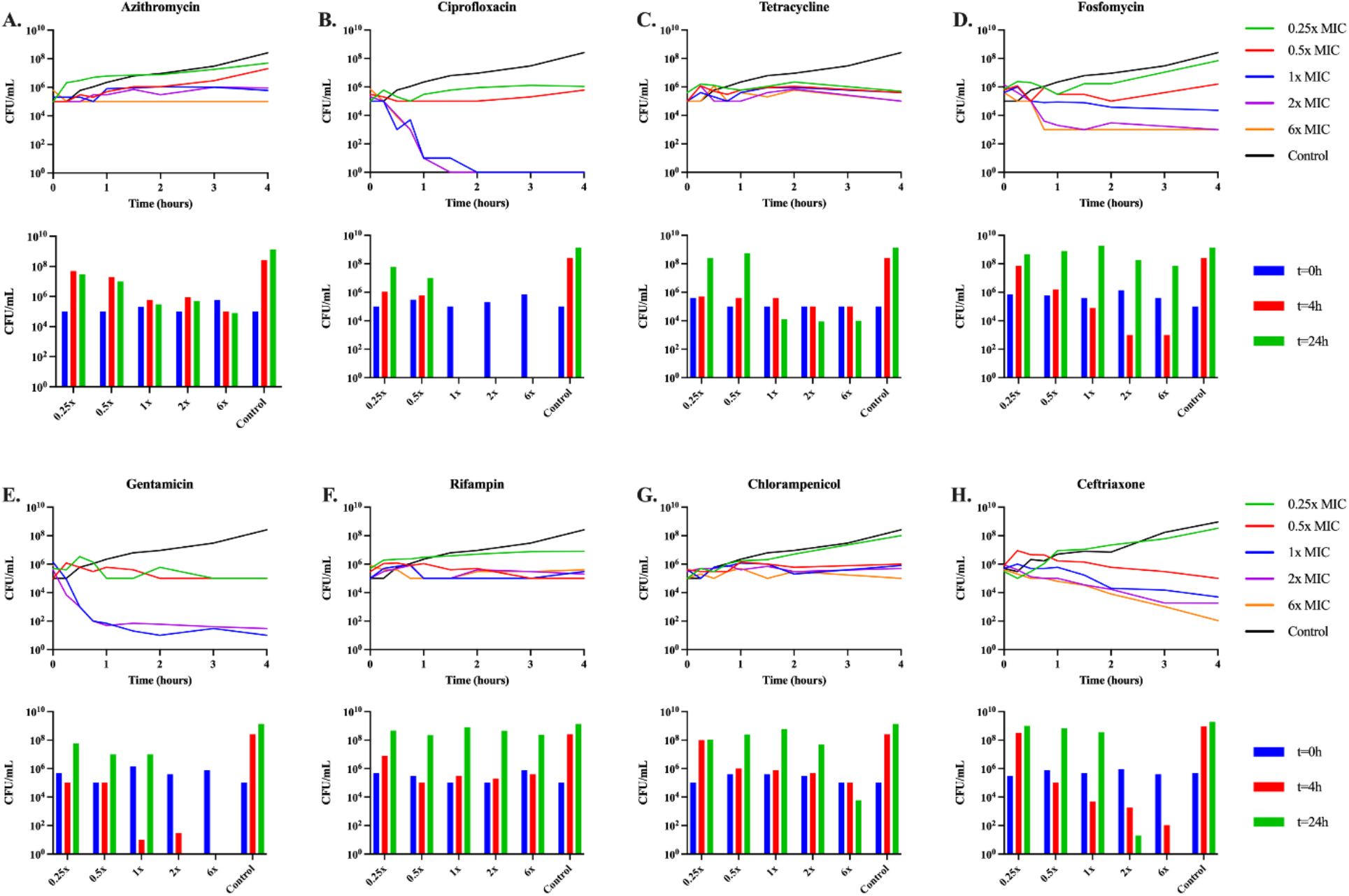
LB time-kills. Changes in viable cell density of E. *coli* MG1655 exposed to sub- and super- MIC concentrations of the eight antibiotics in broth for 24 hours. Shown for each panel in the line graph are changes in the viable cell density over a four-hour period, shown in the bar graph is the viable cell density at 0, 4, and 24 hours. (**A)** Azithromycin (**B)** Ciprofloxacin (**C)** Tetracycline (**D)** Fosfomycin (**E)** Gentamicin (**F)** Rifampin (**G)** Chloramphenicol (**H)** Ceftriaxone.

**Figure 8.**
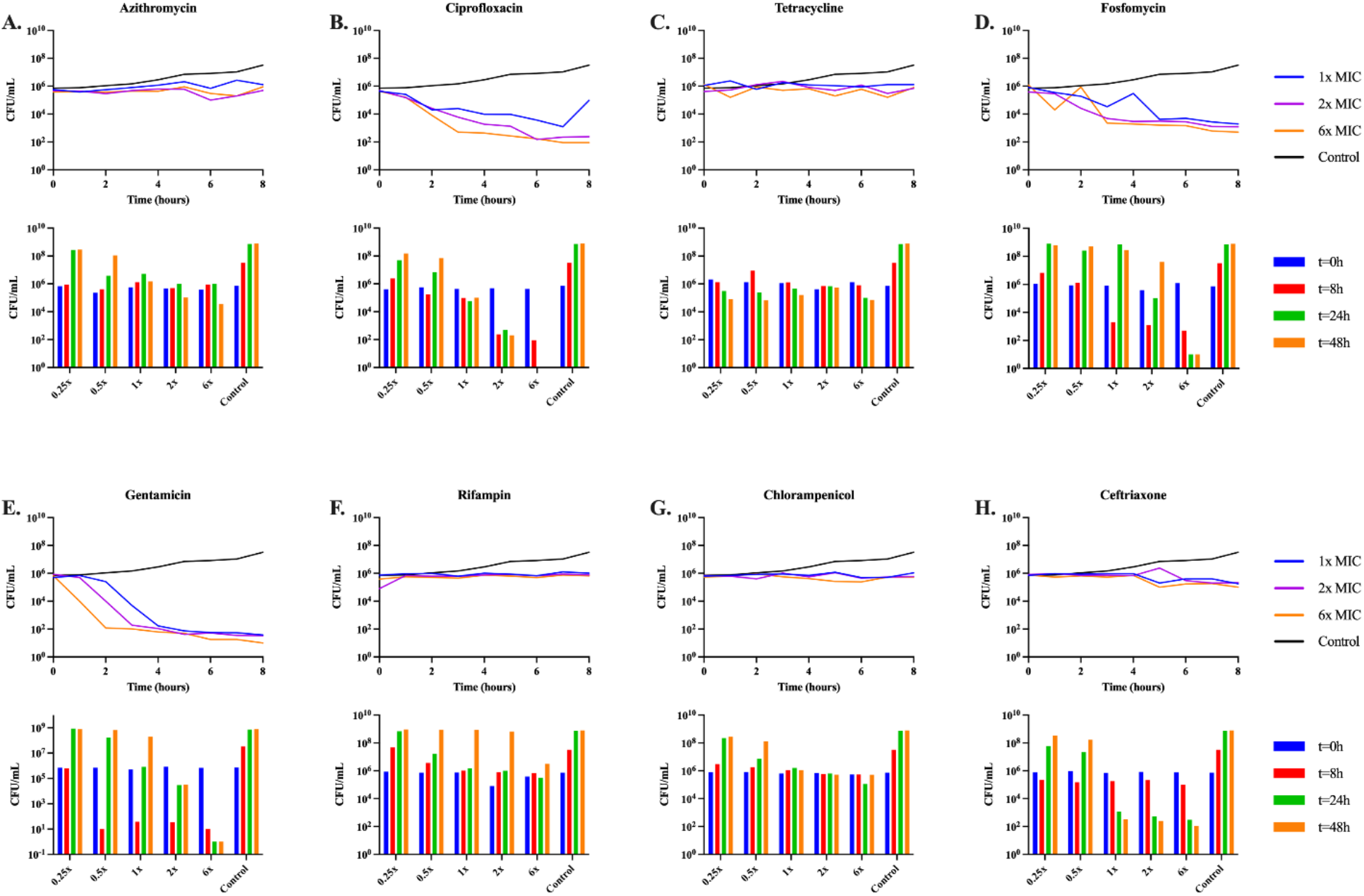
Minimal medium time-kills. Changes in viable cell density of E. *coli* MG1655 exposed to sub- and super- MIC concentrations of the eight antibiotics in minimal for 48 hours. Shown for each panel in the line graph are changes in the viable cell density over a four-hour period, shown in the bar graph is the viable cell density at 0, 8, 24, and 48 hours. Sub-MIC concentrations of the antibiotics are not shown, as over eight hours there was no change in CFUs in any of the cultures. (**A)** Azithromycin (**B)** Ciprofloxacin (**C)** Tetracycline (**D)** Fosfomycin (**E)** Gentamicin (**F)** Rifampin (**G)** Chloramphenicol (**H)** Ceftriaxone.

In both LB and minimal medium, the bacteriostatic drugs (azithromycin, tetracycline, and chloramphenicol) suppress growth of the bacteria at super-MIC concentrations. In both media ciprofloxacin, gentamicin, and ceftriaxone appear highly bactericidal, though ceftriaxone does not kill at all in the first eight hours in minimal media. Rifampin and fosfomycin are both complicated by the emergence of genotypic resistance. Rifampin appears static for the first several hours in both media, but the bacteria end up growing to the level of the control by 24 and 48 hours respectively. Fosfomycin kills the *E. coli* for the first several hours, but by the end of the experiments, the bacterial densities are similar to that of the antibiotic free control.

## Discussion

The diagnosis of diseases and protocols for their treatment are commonly based on relatively few estimated parameters like cholesterol levels, blood pressure, white cell counts, and glucose concentrations. In the case of antibiotic treatment, this parametric reductionism is in the application of the minimum concentration of the antibiotic required to prevent the replication of the bacteria presumed to be responsible for the infection, the MIC. As the unique measure of the pharmacodynamics, MICs are employed as both a measure of the susceptibility of the bacteria to the drug and for the design of dosing regimens for treatment of infections with those bacteria [8, 20]. Under *in vivo* conditions, the local bacterial concentration at the site of infection, the replication rate, the available nutrients, the antibiotic concentration, and the effect of immune response are highly heterogeneous in time and space. In addition, under *in vivo* conditions within tissues, most bacterium are likely exposed to sub-MIC concentrations of drugs derived to an antibiotic gradient [21]. A single, simple test will be insufficient to cover these diverse and changing conditions. Efforts to design culture media mimicking the tissue environment, referred to by Ersoy *et al*. as the “fundamental flaw in the paradigm for antimicrobial susceptibility testing” [22], could slightly improve the *in vitro/in vivo* correlation, but these medias cannot evade the inconvenience of environmental heterogeneity. Our approach is evaluating the effects of nutrients and limiting resources on MIC. Glucose is likely not the main carbon source for *E. coli* in the host [23], but limitations *in vitro* of glucose or other carbon sources mimic the effect of shortage of other possible nutrients. The results of this study with eight antibiotics and *E. coli* illustrate the limitations of MICs for these purposes and also those of more comprehensive measures of the PD of antibiotics, such as Hill functions [24].

MICs are officially estimated with low densities of bacteria introduced into rich media, most commonly broths such as LB or Mueller-Hinton [14, 25]. Our results demonstrate that estimates of this parameter for azithromycin, tetracycline, and gentamicin obtained in glucose-limited minimal medium are markedly lower than those estimated in broth. We have also shown that carbon source plays a large role in the determination of the MIC, particularly for azithromycin and fosfomycin. This is an indication that both nutrient limitation and choice of nutrient plays a large role in the determination of MIC. Our results have also shown a major limitation in the method for estimating MIC. This traditional method inherently cannot account for the emergence of either phenotypic or genotypic resistance.

In addition to MICs, pharmacodynamic Hill functions provide information about the relationship between the concentration of the antibiotic and the dynamics and rates of growth of bacteria exposed to the drug [24]. The parameters of Hill functions are estimated from the antibiotic concentration-dependent growth rates of the bacteria with the assumption being that these drugs only act by modifying these rates and do not take into account changes in any other bacterial growth parameters [24]. The results of this and previous studies [26] question the validity of this assumption.

Our original model, a modification of the Hill functions which allows for resource consumption, predicts that if at sub-MIC concentrations all antibiotics did was to reduce the growth rate of the exposed bacteria, their populations would achieve the same stationary phase densities but do so at times that increase with the concentration of the antibiotic. To an extent that varies among drugs, sub-MIC concentrations of antibiotics increase the time before the bacteria start to grow, the lag. This effect is most prominent among drugs that alter cellular metabolism. Antibiotics slowing metabolic processes likely slow development of cellular structures, as all structural elements of the cell will derive from glucose metabolism, which would be seen as an increase in the lag time. Also unanticipated by our model extending the Hill functions is that the maximum density to which the bacteria grow is dependent on the antibiotic concentration. Though, our updated model, which varies the conversion efficiency of glucose with an increasing concentration of an antibiotic, can account for these effects.

One possible explanation for why sub-MIC concentrations of antibiotics lowers the densities to which the exposed populations of bacteria grow is that these drugs reduce the efficacy of use of the limiting resource. Stated another way, the bacteria exposed to sub-inhibitory antibiotic concentrations require more resource for replication. The effect of antibiotics on bacterial nutritional efficiency can explain this result. Certainly, antibiotic exposure decreases the metabolic flux, in that there is a waste of intracellular metabolites that are synthesized but not integrated into metabolic pathways. For instance, exposure to chloramphenicol and rifampin results in an accumulation of amino acids, nucleotides, and lipids [27]. Similarly, beta-lactam exposure induces a futile cycle of cell wall synthesis and degradation, which depletes cellular resources [28]. There is also emerging evidence that sub-MIC concentrations of antibiotics can influence non-essential tRNA and RNA modifications, leading to unnecessary energy expenditures [29]. Such energetic waste should be compensated by the consumption of nutrients that do not necessarily contribute to the net growth rate.

Consistent with this hypothesis at sub-MIC concentrations, all of the antibiotic treated cultures fail to grow up to the glucose-limited density of the controls, but yet all of the glucose is utilized in that no or very little free glucose is detectable. While, at super-MIC concentrations the bacterial densities are suppressed, and a substantial amount of free glucose remains. As a departure from this trend, the cultures treated with super-MIC concentrations of ceftriaxone use more free glucose than the cultures treated with super-MIC concentrations of the other drugs, but yet a large amount of glucose is still present. It has not escaped our notice that based on Figure 8 this beta-lactam over the first eight hours appears very similar to the bacteriostatic drugs. Could it be that in these first few hours, the growth rate is equal to the death rate so that these bacteria are still utilizing the free glucose? Alternatively, ceftriaxone is known to produce filaments due to PBP3 inhibition, could it be that these multicellular filaments have an increased metabolic requirement when compared to their ancestor cells?

Advancement in the way we measure and understand pharmacodynamics should have therapeutic implications. The most used PD parameter, the MIC, certainly allows for the comparison of antibiotic susceptibility between two isolates of the same species under identical *in vitro* conditions. However, MIC is an insufficient PD marker for several reasons. Firstly, the antibiotic effect at subinhibitory concentrations is not considered. Secondly, this parameter does not tell us anything about the susceptibility of a given organism at different cell densities. Thirdly, it tells us nothing about changes in growth parameters such as the effect on lag, average growth rate, final density, or overall rate of drug-mediated killing. Lastly, MIC indicates nothing about post-antibiotic effects, resistance, persistence, or heteroresistance. All these pharmacodynamic dimensions change the effect of antibiotics *in vivo* as the concentration of the drug changes over time, as predicted by the PK. Without understanding these effects, the operative linkage of PK/PD is insufficient to obtain robust predictions about antibiotic efficacy. When MIC is used as the sole PD parameter, this insufficiency is even more apparent. Most PK/PD approaches look at one of three PK parameters: either maximum concentration of the antibiotic to MIC, drug exposure to MIC (which is to say area under the curve to MIC), or the time that the drug concentration is above the MIC [30]. Most importantly, the development of novel antibiotics should consider these PD complexities and adopt a non-reductive approach which is not simply limited to MIC. An antibiotic should be more effective than another as shown by a holistic superiority-test, considering multiple parameters for PD. These tests should take into account: the effect of the density of the starting bacterial population, the effectiveness of the drug on bacteria in different stages of growth, the ability to reduce bacterial growth rate, increase lag time, and decrease final population size, as well as decrease the number of persisters. Optimally, all these advantages would also reduce the length of treatment, the emergence of antibiotic resistance, and the recurrent infections, as well as the possibility of transmission of pathogens to other hosts. However, the usual microbiological tests in drug development do not include these calculations, benefits which would go unrecognized by using an MIC-focused approach.

## Materials and Methods

### Numerical solutions (simulations)

For our numerical analysis of the equations presented (equations 1–6), we used Berkeley Madonna, using parameters in the ranges estimated for *E. coli*. Copies of the Berkeley Madonna program used for these simulations are available at www.eclf.net.

### Growth media

LB broth (244620) was obtained from BD. The DM (Davis Minimal) minimal base without dextrose (15758-500G-F) was obtained from Sigma Aldrich. Glucose (41095-5000) was obtained from Acros, Succinic Acid (S-2378) was obtained from Sigma Aldrich, Lactose (L3750) was obtained from Sigma Aldrich, Maltose (M2250) was obtained from Sigma Aldrich, Glycerol (G7757) was obtained from Honeywell and Fructose (161355000) was obtained from ACROS Organics. LB agar (244510) for plates was obtained from BD.

### Bacterial strains

*E. coli* MG1655 was obtained from the Levin Lab Bacterial collection.

### Antibiotics

Ciprofloxacin (A4556) was obtained from AppliChem Panreac, Tetracycline (T17000) was obtained from Research Products International, Fosfomycin (P5396) was obtained from Sigma Aldrich, Chloramphenicol (23660) was obtained from USB, Rifampin (BP2679-1) was obtained from Fisher BioReagents, Ceftriaxone (C5793) was obtained from Sigma Aldrich, Azithromycin (3771) was obtained from TOCRIS and Gentamicin (BP918-1) was obtained from Fisher BioReagents.

### Sampling bacterial densities

The densities of bacteria were estimated by serial dilution in 0.85% saline and plating. The total density of bacteria was estimated on Lysogeny Broth (LB) plates with 1.6% agar.

### Growth rate estimation

Exponential growth rates were estimated from changes in optical density (OD600) in a Bioscreen C. For this, 24-hours stationary phase cultures were diluted in LB or glucose-limited liquid media to an initial density of approximately 10^5^ cells per mL. Five replicas were made for each estimate by adding 300μl of the suspensions to the wells of the Bioscreen plates. The plates were incubated at 37°C and shaken continuously. Estimates of the OD (600nm) were made every five minutes for 24 hours in LB and 48 hours in glucose-limited medium. Normalization, replicate means and error, growth rate, lag and maximum OD were found using a novel R Bioscreen C analysis tool accessible at https://josheclf.shinyapps.io/bioscreen_app.

### Sequencing and analyses

Bacterial DNA was extracted using Promega’s (Wisconsin, USA) Wizard Genomics DNA Purification Kit (Cat# A1120) using the manufacturer’s protocol. The extracted DNA was quantified on ThermoFisher’s NanoDrop OneC microvolume spectrophotometer (Cat# ND-ONEW). Samples were sent to the Microbial Genome Sequencing Center in Pittsburgh, Pennsylvania, USA, for whole genome sequencing on the Illumina NextSeq 2000 platform. Analysis of FASTAq files received from the Microbial Genome Sequencing Center were analyzed using Geneious Prime version 2022.0.1.

### Minimum inhibitory concentrations

Antibiotic MICs were estimated using a 2-fold microdilution procedure as described in [31].

### Antibiotic time-kills

For these experiments, overnight cultures of *E. coli* MG1655 were added to LB broth or DM 1000 μg/mL glucose liquid media at an initial density of approximately 10^5^ cells per mL, followed by 1 h incubation at 37°C and shaken continuously. After incubation, antibiotics were added at the concentrations indicated in Figures 4 and 5 and the initial densities of the cultures were estimated. The cultures with the antibiotics were incubated for 24 h in LB media and 48 h in DM glucose and estimates of the viable cell densities were made at 15 min, 30 min, 45 min, 1 h, 1.5 h, 2 h, 3 h, 4 h and 24 h for LB cultures and 1 h, 2 h, 3 h, 4 h, 5 h, 6 h, 7 h, 8 h, 24 h and 48 h for the DM glucose liquid media cultures.

### Glucose assay

Glucose concentration was measured by a dinitrosalicylic colorimetric method as described in [32].

## Acknowledgments

We would like to thank Danielle Steed, Michael Woodworth, and Ahmed Babiker for their discussions of the clinical implications and relevance of this work. We would also like to thank Nic Vega, Ingrid McCall, and the other members of the Levin Lab.

BRL would like to thank the US National Institute of General Medical Sciences for their funding support via R35 GM 136407 and the US National Institute of Allergy and Infectious Diseases for their funding support via U19 AI 158080-02.

TGG would like to thank the US Fulbright Program and the Spanish Government FPI fellowship for their funding support.

## Data Availability Statement

All the data generated are available in this manuscript and its supporting supplemental material.

## Supplementary Material

**Fig. S1.**
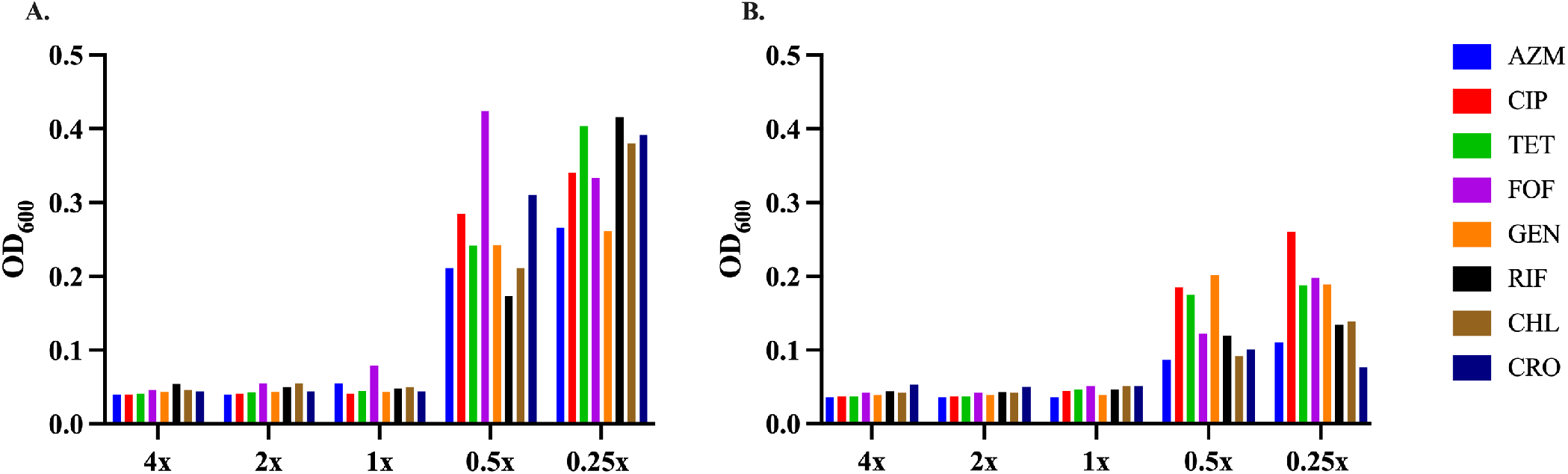
MIC ooze. Antibiotic MICs were performed using a 2-fold microdilution procedure and after 24 hours in **(A)** LB and 48 h in **(B)** glucose-limited minimal medium the final optical density (OD600nm) was measured at different concentrations of antibiotic.

**Fig. S2.**
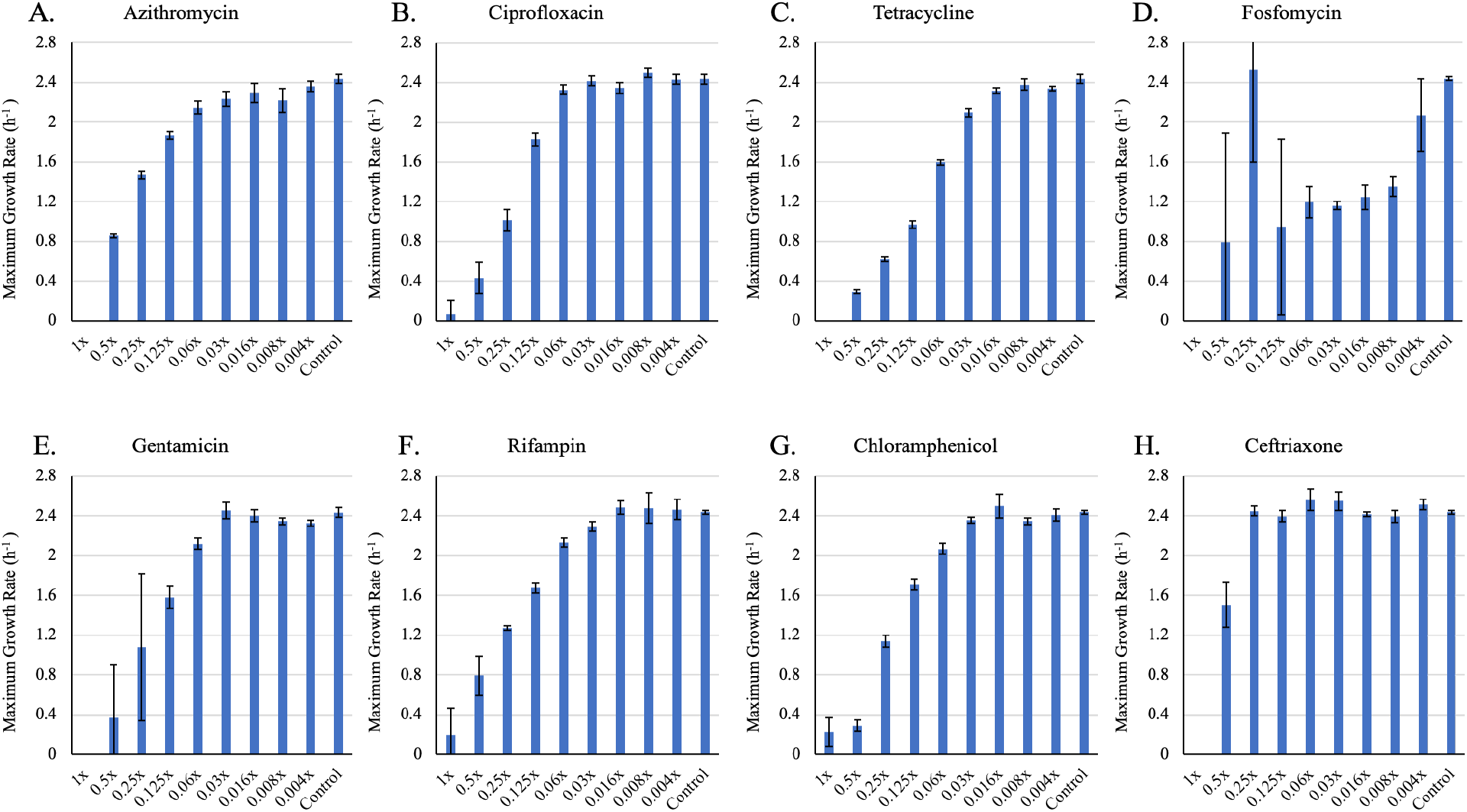
**Changes in maximum growth rate (*v_MAX_*)** of E. *coli* MG1655 exposed to different sub-MIC concentrations of eight antibiotics for 24 hours in LB. Bars are representative of the average of five technical replicas. Each concentration is shown as a fraction of the MIC for the noted drug (**A)** Azithromycin (**B)** Ciprofloxacin (**C)** Tetracycline (**D)** Fosfomycin (**E)** Gentamycin (**F)** Rifampin (**G)** Chloramphenicol (**H)** Ceftriaxone.

**Fig. S3.**
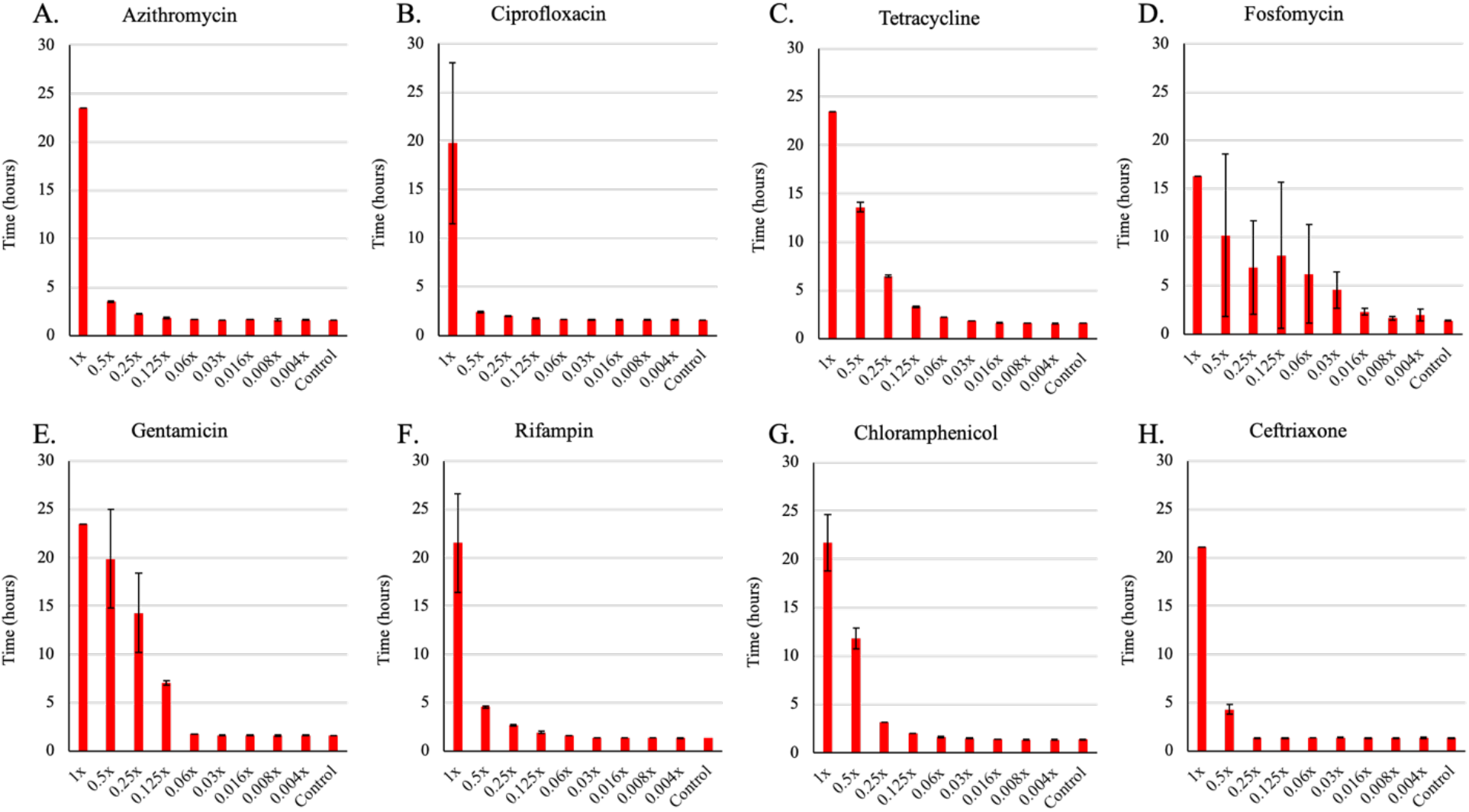
**Changes in the time before the bacteria start to grow (lag)** of E. *coli* MG1655 exposed to different sub-MIC concentrations of eight antibiotics for 24 hours in LB. Bars are representative of the average of five technical replicas. Each concentration is shown as a fraction of the MIC for the noted drug (**A)** Azithromycin (**B)** Ciprofloxacin (**C)** Tetracycline (**D)** Fosfomycin (**E)**

**Fig. S4.**
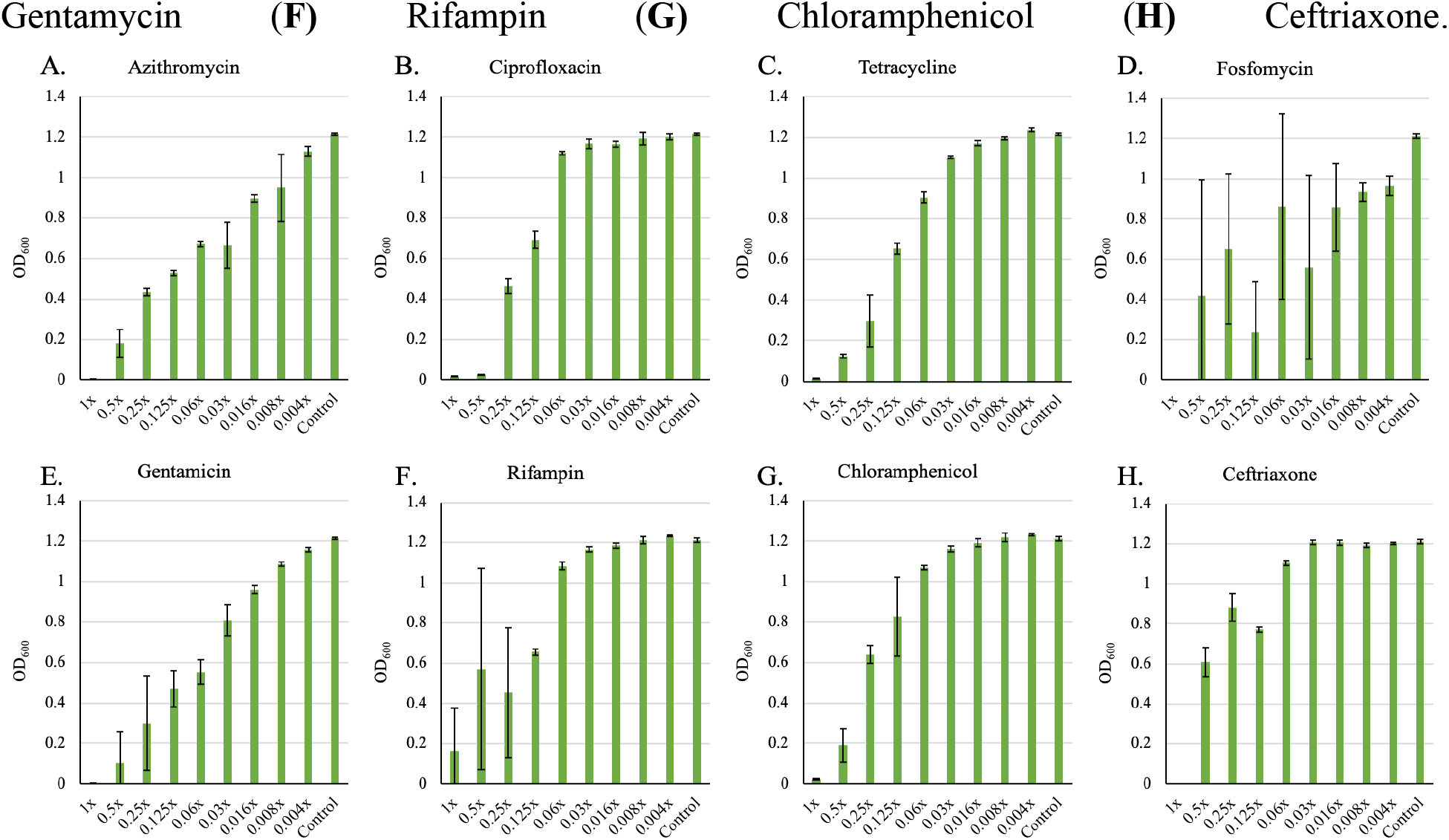
**Changes in the maximum optical density (OD 600nm)** of E. *coli* MG1655 exposed to different sub-MIC concentrations of eight antibiotics for 24 hours in LB. Bars are representative of the average of five technical replicas. Each concentration is shown as a fraction of the MIC for the noted drug (**A)** Azithromycin (**B)** Ciprofloxacin (**C)** Tetracycline (**D)** Fosfomycin (**E)** Gentamycin (**F)** Rifampin (**G)** Chloramphenicol (**H)** Ceftriaxone.

**Fig. S5.**
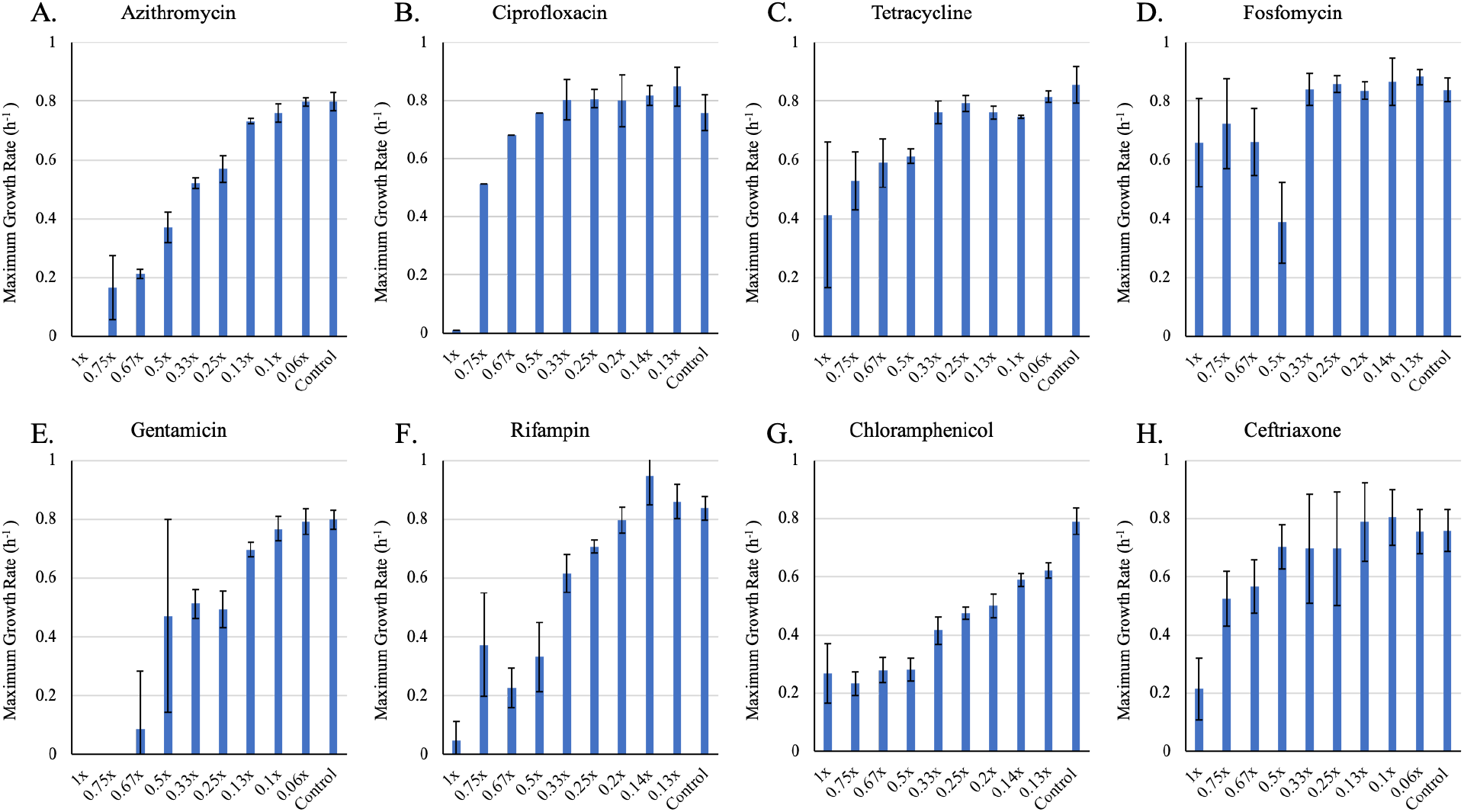
**Changes in maximum growth rate (*v_MAX_*)** of E. *coli* MG1655 exposed to different sub-MIC concentrations of eight antibiotics for 48 hours in glucose-limited minimal medium. Bars are representative of the average of five technical replicas. Each concentration is shown as a fraction of the MIC for the noted drug (**A)** Azithromycin (**B)** Ciprofloxacin (**C)** Tetracycline (**D)** Fosfomycin (**E)** Gentamycin (**F)** Rifampin (**G)** Chloramphenicol (**H)** Ceftriaxone.

**Fig. S6.**
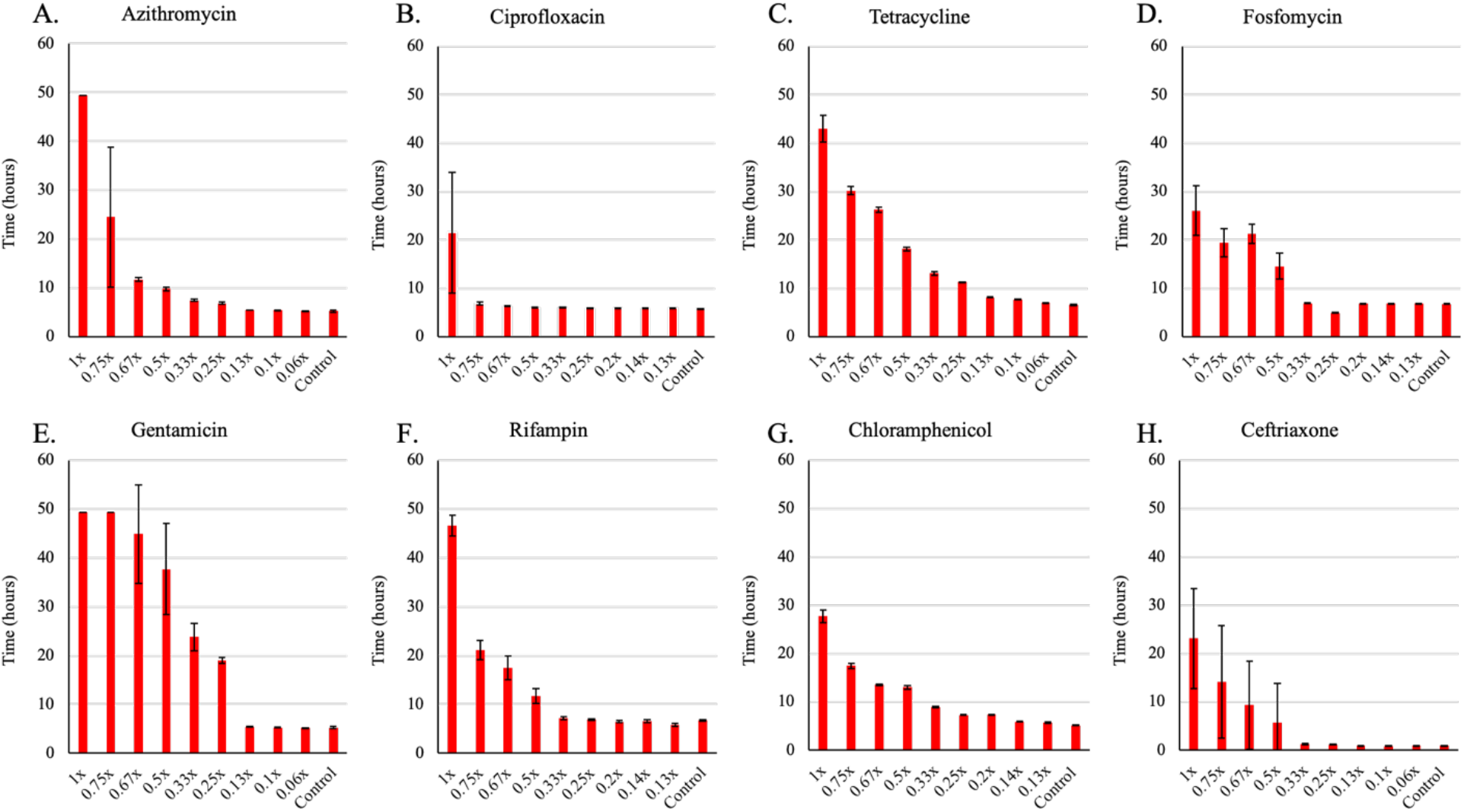
**Changes in the time before the bacteria start to grow (lag)** of E. *coli* MG1655 exposed to different sub-MIC concentrations of eight antibiotics for 48 hours in glucose-limited minimal medium. Bars are representative of the average of five technical replicas. Each concentration is shown as a fraction of the MIC for the noted drug (**A)** Azithromycin (**B)** Ciprofloxacin (**C)** Tetracycline (**D)** Fosfomycin (**E)** Gentamycin (**F)** Rifampin (**G)** Chloramphenicol (**H)** Ceftriaxone.

**Fig. S7.**
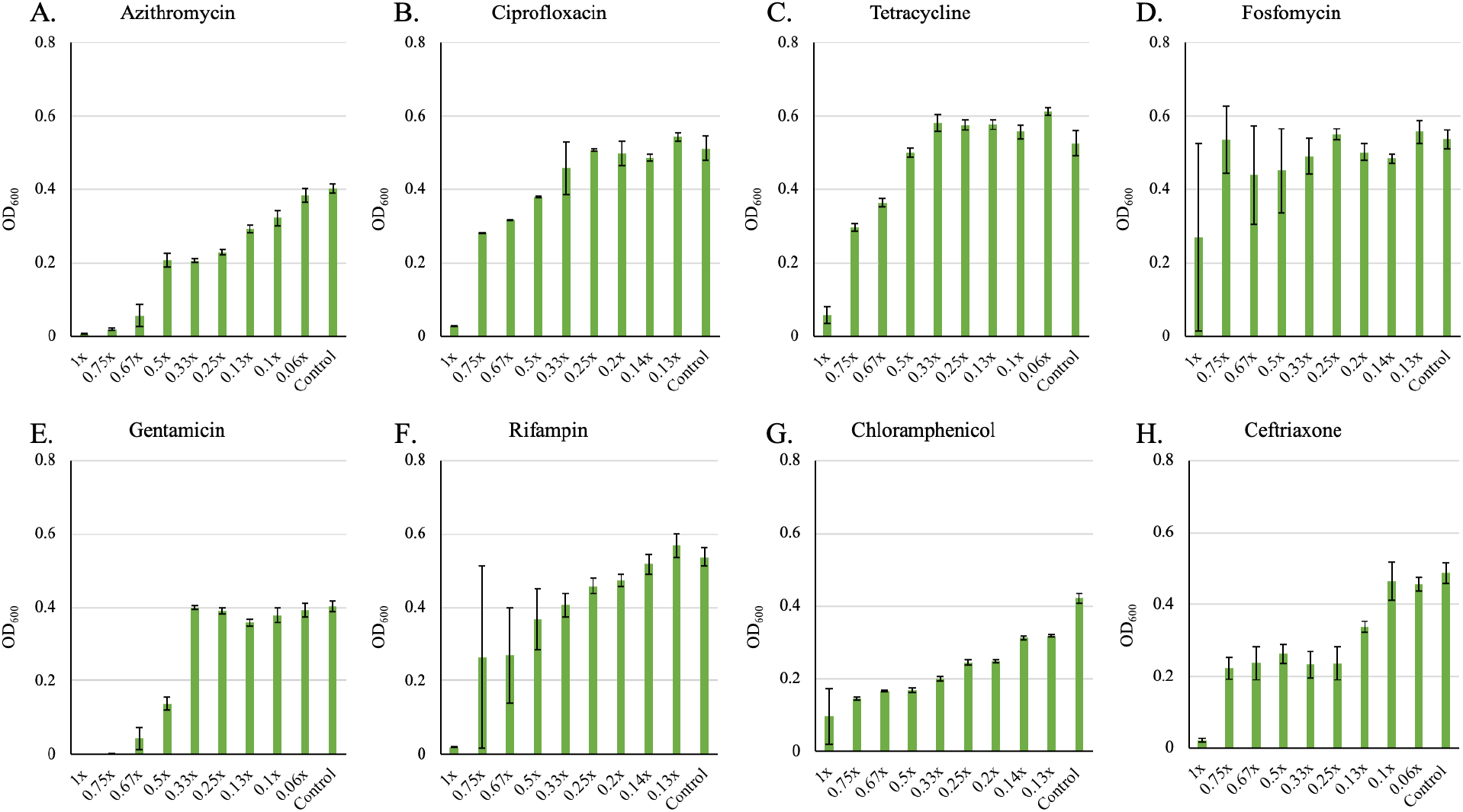
**Changes in the maximum optical density (OD 600nm)** of E. *coli* MG1655 exposed to different sub-MIC concentrations of eight antibiotics for 48 hours in glucose-limited minimal medium. Bars are representative of the average of five technical replicas. Each concentration is shown as a fraction of the MIC for the noted drug (**A)** Azithromycin (**B)** Ciprofloxacin (**C)** Tetracycline (**D)** Fosfomycin (**E)** Gentamycin (**F)** Rifampin (**G)** Chloramphenicol (**H)** Ceftriaxone.

**Fig. S8.**
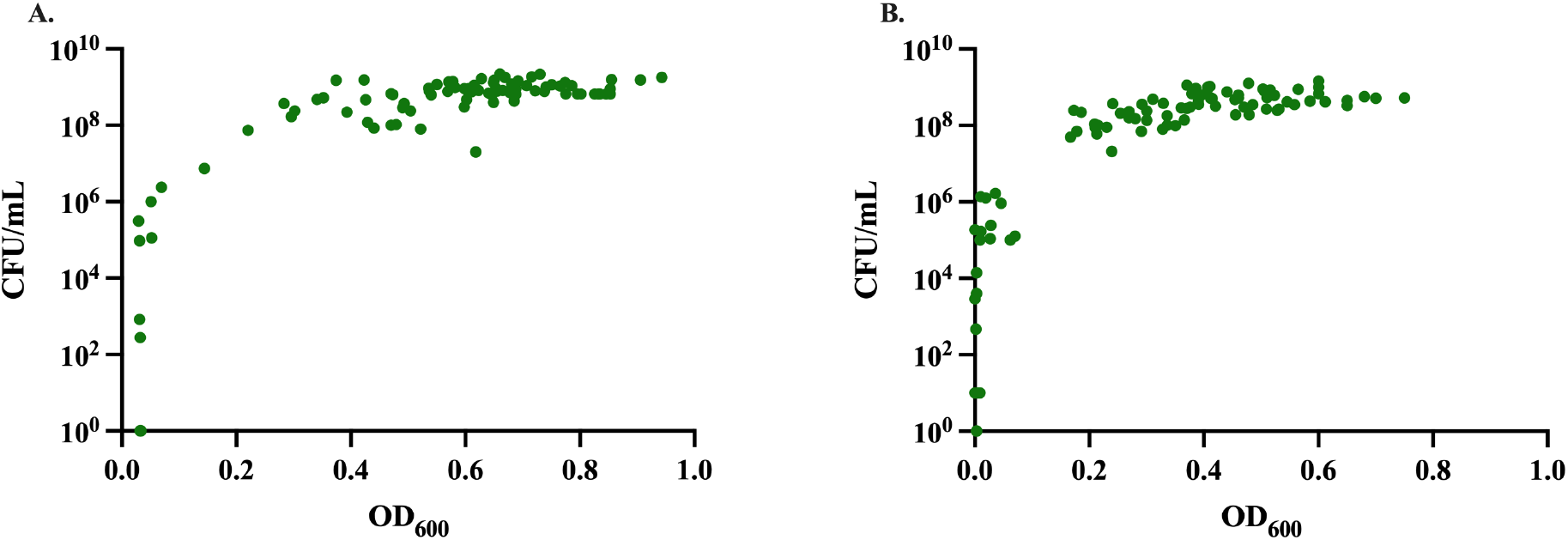
**Relation between optical density and viable colony forming units (CFU)** of E. *coli* MG1655 cultures exposed to sub- and super- MIC concentrations of the eight antibiotics in **(A)** LB after 24 hours and in **(B)** glucose-limited medium after 48 hours of incubation.

**Figure S9.**
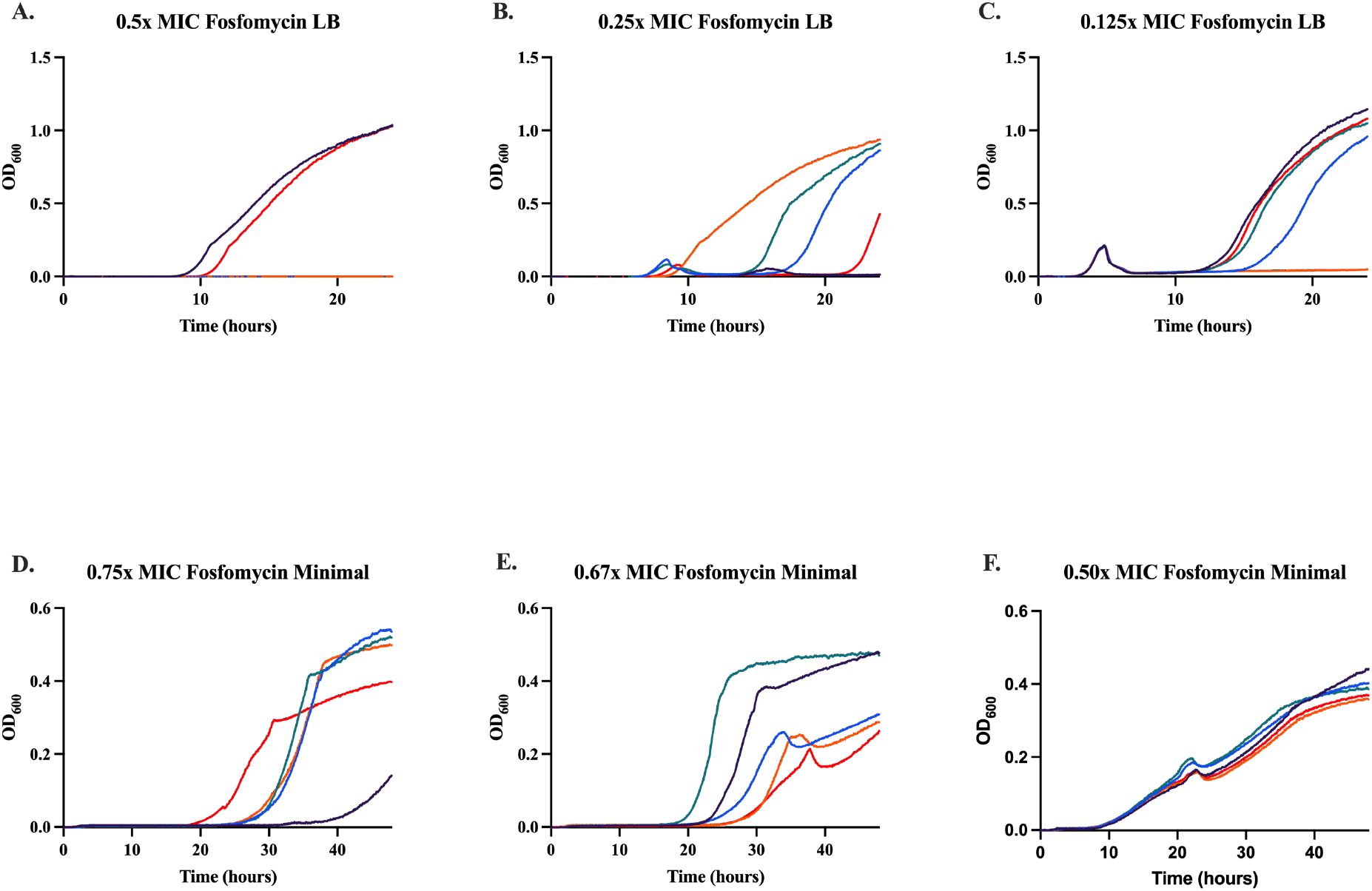
Growth dynamics in LB and glucose-limited medium at different fosfomycin concentrations. Changes in optical density (600nm) of *E. coli* MG1655 exposed to different fosfomycin concentrations for 24 hours in LB and 48 h in glucose-limited medium. Lines are representative of one technical replica and normalized to the time zero optical density. Individual replicate curves show how resistance came up at different fosfomycin concentrations.

**Figure S10.**
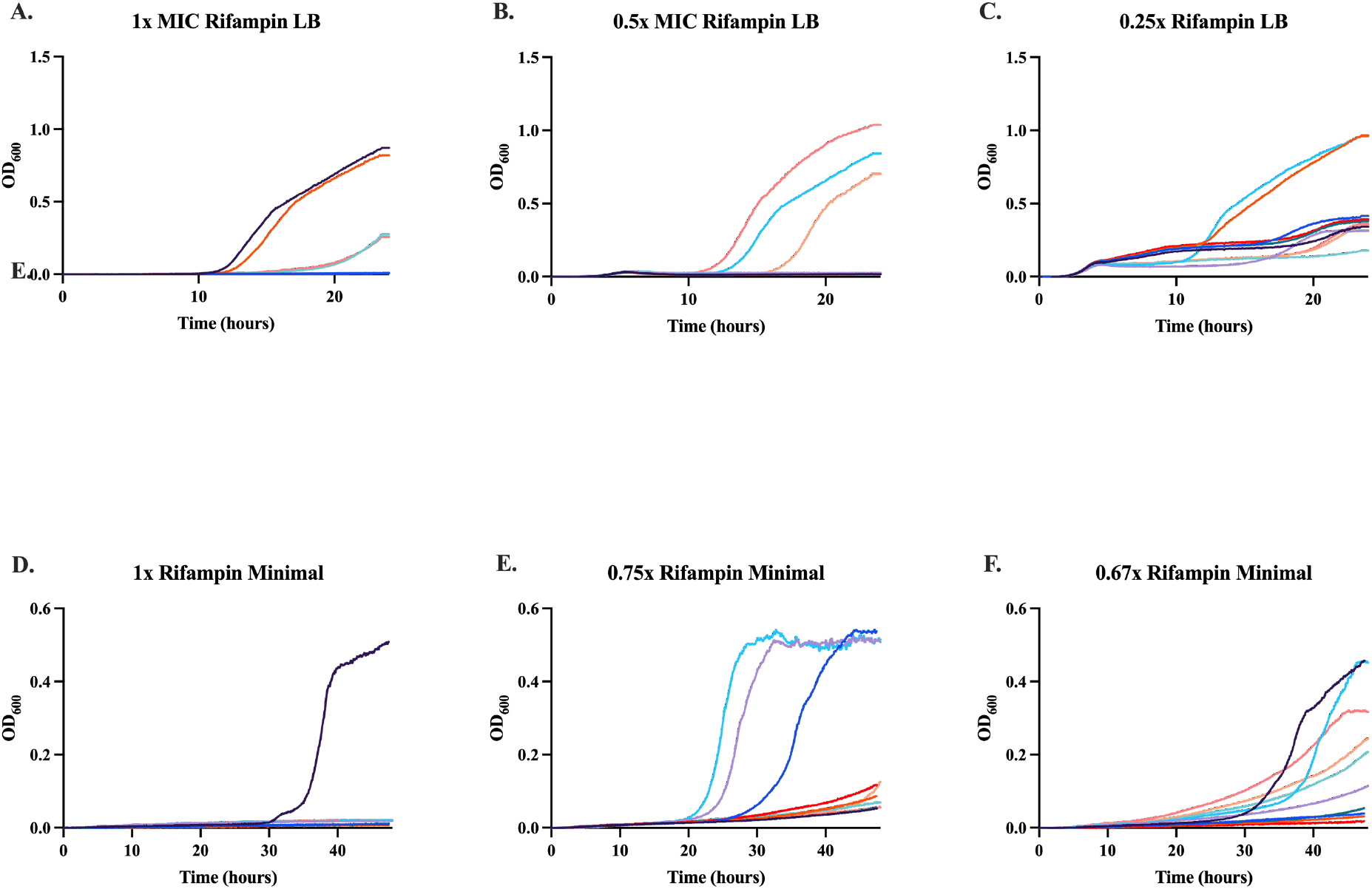
Growth dynamics in LB and glucose-limited medium at different rifampin concentrations. Changes in optical density (600nm) of *E. coli* MG1655 exposed to different rifampin concentrations for 24 hours in LB and 48 h in glucose-limited medium. Lines are representative of one technical replica and normalized to the time zero optical density. Individual replicate curves show how resistance came up at different rifampin concentrations.

**Table S1.**
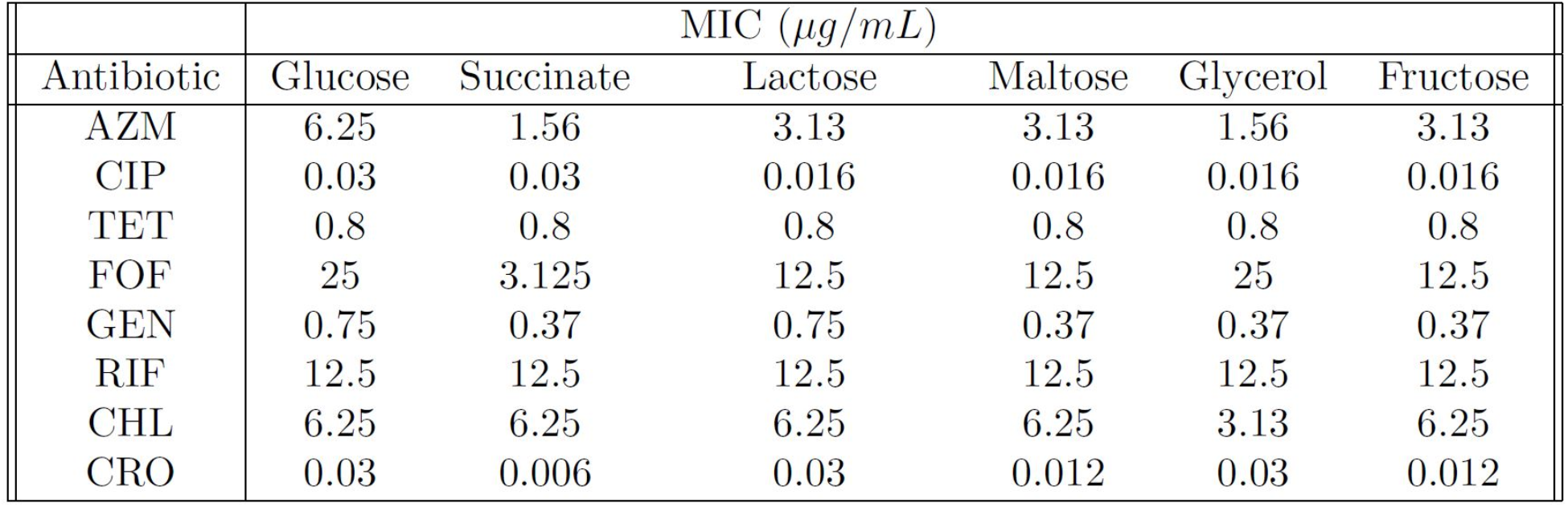
Estimated MICs of *E. coli* MG1655 for eight antibiotics in limited minimal medium with different carbon sources.

## References

1. Andes, D., et al., Application of pharmacokinetics and pharmacodynamics to antimicrobial therapy of respiratory tract infections. Clin Lab Med, 2004. 24(2): p. 477–502.

2. Craig, W.A. and D. Andes, Pharmacokinetics and pharmacodynamics of antibiotics in otitis media. Pediatric Infectious Disease Journal, 1996. 15(3): p. 255–9.

3. Velkov, T., et al., PK/PD models in antibacterial development. Curr Opin Microbiol, 2013. 16(5): p. 573–9.

4. Frimodt-Moller, N., How predictive is PK/PD for antibacterial agents? Int J Antimicrob Agents, 2002. 19(4): p. 333–9.

5. Levison, M.E. and J.H. Levison, Pharmacokinetics and pharmacodynamics of antibacterial agents. Infect Dis Clin North Am, 2009. 23(4): p. 791–815, vii.

6. Rawson, T.M., et al., Exploring the Pharmacokinetics of Phenoxymethylpenicillin (Penicillin-V) in Adults: A Healthy Volunteer Study. Open Forum Infectious Diseases, 2021. 8(12).

7. Khan, D.D., L.E. Friberg, and E.I. Nielsen, A pharmacokinetic–pharmacodynamic (PKPD) model based on in vitro time–kill data predicts the in vivo PK/PD index of colistin. Journal of Antimicrobial Chemotherapy, 2016. 71(7): p. 1881–1884.

8. Landersdorfer, C.B. and R.L. Nation, Limitations of Antibiotic MIC-Based PK-PD Metrics: Looking Back to Move Forward. Front Pharmacol, 2021. 12: p. 770518.

9. Lorian, V., Antibiotics in Laboratory Medicine. 2005: Lippincott Williams & Wilkins.

10. Kowalska-Krochmal, B. and R. Dudek-Wicher, The Minimum Inhibitory Concentration of Antibiotics: Methods, Interpretation, Clinical Relevance. Pathogens, 2021. 10(2).

11. Wikler, M.A., Methods for dilution antimicrobial susceptibility tests for bacteria that grow aerobically: approved standard. Clsi (Nccls), 2006. 26: p. M7–A7.

12. Ling, T.K., Z.K. Liu, and A.F. Cheng, Evaluation of the VITEK 2 system for rapid direct identification and susceptibility testing of gram-negative bacilli from positive blood cultures. J Clin Microbiol, 2003. 41(10): p. 4705–7.

13. Bobenchik, A.M., et al., Performance of Vitek 2 for antimicrobial susceptibility testing of Enterobacteriaceae with Vitek 2 (2009 FDA) and 2014 CLSI breakpoints. J Clin Microbiol, 2015. 53(3): p. 816–23.

14. Colombo, A.L., et al., Comparison of Etest and National Committee for Clinical Laboratory Standards broth macrodilution method for azole antifungal susceptibility testing. J Clin Microbiol, 1995. 33(3): p. 535–40.

15. Amsler, K., et al., Comparison of Broth Microdilution, Agar Dilution, and Etest for Susceptibility Testing of Doripenem against Gram-Negative and Gram-Positive Pathogens. Journal of Clinical Microbiology, 2010. 48(9): p. 3353–3357.

16. Udekwu, K.I., et al., Functional relationship between bacterial cell density and the efficacy of antibiotics. J Antimicrob Chemother, 2009. 63(4): p. 745–57.

17. Stokes, J.M., et al., Bacterial Metabolism and Antibiotic Efficacy. Cell Metab, 2019. 30(2): p. 251–259.

18. Monod, J., THE GROWTH OF BACTERIAL CULTURES. Annual Review of Microbiology, 1949. 3(1): p. 371–394.

19. Ballestero-Téllez, M., et al., Role of inoculum and mutant frequency on fosfomycin MIC discrepancies by agar dilution and broth microdilution methods in Enterobacteriaceae. Clinical Microbiology and Infection, 2017. 23(5): p. 325–331.

20. Benkova, M., O. Soukup, and J. Marek, Antimicrobial susceptibility testing: currently used methods and devices and the near future in clinical practice. Journal of Applied Microbiology, 2020. 129(4): p. 806–822.

21. Baquero, F. and M.C. Negri, Selective compartments for resistant microorganisms in antibiotic gradients. Bioessays, 1997. 19(8): p. 731–6.

22. Ersoy, S.C., et al., Correcting a Fundamental Flaw in the Paradigm for Antimicrobial Susceptibility Testing. EBioMedicine, 2017. 20: p. 173–181.

23. Brown, S.A., K.L. Palmer, and M. Whiteley, Revisiting the host as a growth medium. Nat Rev Microbiol, 2008. 6(9): p. 657–66.

24. Regoes, R.R., et al., Pharmacodynamic functions: a multiparameter approach to the design of antibiotic treatment regimens. Antimicrob Agents Chemother, 2004. 48(10): p. 3670–6.

25. Leclercq, R., et al., EUCAST expert rules in antimicrobial susceptibility testing. Clin Microbiol Infect, 2013. 19(2): p. 141–60.

26. Atkinson, B.A., L. Amaral, and J.A. Washington, Sublethal Concentrations of Antibiotics, Effects on Bacteria and the Immune System. CRC Critical Reviews in Microbiology, 1982. 9(2): p. 101–138.

27. Lobritz, M.A., et al., Antibiotic efficacy is linked to bacterial cellular respiration. Proc Natl Acad Sci U S A, 2015. 112(27): p. 8173–80.

28. Cho, H., T. Uehara, and T.G. Bernhardt, Beta-lactam antibiotics induce a lethal malfunctioning of the bacterial cell wall synthesis machinery. Cell, 2014. 159(6): p. 1300–11.

29. Babosan, A., et al., Non-essential tRNA and rRNA modifications impact the bacterial response to sub-MIC antibiotic stress. bioRxiv, 2022: p. 2022.02.06.479318.

30. Eyler, R.F. and K. Shvets, Clinical Pharmacology of Antibiotics. Clin J Am Soc Nephrol, 2019. 14(7): p. 1080–1090.

31. Jorgensen, J.H. and M.J. Ferraro, Antimicrobial susceptibility testing: a review of general principles and contemporary practices. Clin Infect Dis, 2009. 49(11): p. 1749–55.

32. Miller, G.L., Use of dinitrosalicylic acid reagent for determination of reducing sugar. Analytical chemistry, 1959. 31(3): p. 426–428.

